# PRMT5 in T cells drives Th17 responses, mixed granulocytic inflammation and severe allergic airway inflammation

**DOI:** 10.1101/2021.10.13.464281

**Authors:** Brandon W. Lewis, Stephanie A. Amici, Hye-Young Kim, Emily Shalosky, Aiman Khan, Joshua Walum, Kymberly M. Gowdy, Joshua A. Englert, Ned A. Porter, Mitchell H. Grayson, Rodney D. Britt, Mireia Guerau-de-Arellano

**Affiliations:** Center for Perinatal Research, The Abigail Wexner Research Institute at Nationwide Children’s Hospital, Columbus, OH; Division of Medical Laboratory Science, Wexner Medical Center, School of Health and Rehabilitation Sciences, Columbus, OH; Department of Chemistry and Vanderbilt Institute of Chemical Biology, Vanderbilt University, Nashville, TN; Division of Pulmonary, Critical Care and Sleep Medicine, The Ohio State University Wexner Medical Center, Davis Heart and Lung Research Institute, Columbus, OH; Center for Clinical and Translational Research, The Abigail Wexner Research Institute at Nationwide Children’s Hospital, Columbus, OH; Division of Allergy and Immunology, The Ohio State University, Columbus, OH; Department of Pediatrics, The Ohio State University Wexner Medical Center, The Ohio State University, Columbus, OH; Institute for Behavioral Medicine Research, The Ohio State University, Columbus, OH; Departments of Microbial Infection and Immunity, The Ohio State University, Columbus, OH; Neuroscience, The Ohio State University, Columbus, OH

## Abstract

Severe asthma is characterized by steroid insensitivity and poor symptom control, and is responsible for the majority of asthma-related hospital costs. Therapeutic options remain limited, in part due to limited understanding in mechanisms driving severe asthma. Increased arginine methylation, catalyzed by protein arginine methyltransferases (PRMTs), is increased in asthmatic lungs. Here, we show that PRMT5 drives allergic airway inflammation in a mouse model reproducing multiple aspects of human severe asthma. We find that PRMT5 is required in CD4^+^ T cells for chronic steroid-insensitive severe lung inflammation, with selective T cell deletion of PRMT5 robustly suppressing eosinophilic and neutrophilic lung inflammation, pathology, airway remodeling and hyperresponsiveness. Mechanistically, we observed high pulmonary sterol metabolic activity, ROR-γt and Th17 responses, with PRMT5-dependent increases in ROR-γt’s agonist desmosterol. Our work demonstrates that T cell PRMT5 drives severe allergic lung inflammation and has potential implications for the pathogenesis and therapeutic targeting of severe asthma.

## Introduction

Asthma is a heterogenous inflammatory lung disease characterized by reversible airflow obstruction and pulmonary inflammation that affects more than 24 million in the US and approximately 238 million people worldwide (1), with an estimated cost of $56 billion (2, 3). It’s esztimated that 40-70% of asthma hospital costs originate from severe asthmatics, who experience greater disease severity, more frequent exacerbations, and are insensitive to corticosteroid treatment (2–6). Phenotypically, severe asthmatics with neutrophilic or mixed neutrophilic/eosinophilic lung infiltrates are less likely to benefit from corticosteroids or therapies that solely target the eosinophilic component of asthma (7–9). Therefore, understanding mechanisms driving mixed granulocytic infiltration and severe asthma could provide opportunities to develop novel therapies for severe asthma.

Protein arginine methylation is an important post-translational modification that regulates signal transduction, DNA repair, RNA processing, protein-protein interactions and gene expression (10, 11). Protein arginine methyl transferases (PRMTs) are classified into type I, II or III enzymes based on their ability to catalyze asymmetric dimethylation (ADM), symmetric dimethylation (SDM) or monomethylation of target proteins (11). Among PRMTs, PRMT1 and PRMT5 are responsible for the majority of ADM and SDM, respectively, in the cell. Enhanced arginine methylation levels have been found in allergen-challenged mice, as well as in lung and sputum samples of asthma patients (12, 13). However, whether arginine methylation plays a role in allergic airway disease, as well as whether PRMTs act through immune or structural cells, remains unknown.

One of the barriers to understanding and treating severe asthma is the difficulty of reproducing multiple phenotypes of severe asthma in animal models of allergic airway inflammation. These severe asthma characteristics include chronicity, neutrophilic or mixed granulocytic infiltration, steroid insensitivity, airway remodeling and airway hyperreactivity (AHR). Short-term exposure of mice to intranasal c-di-GMP (GMP) and house dust mite (HDM) allergen exposure provides one of the first models of severe asthma, which was characterized by eosninophil/neutrophil infiltration, steroid-resistant airway inflammation and airway hyperresponsiveness (AHR) (14). Such models may help uncover mechanisms driving severe asthma and test the effectiveness of novel drugs on established disease.

Here, we find that using the novel GMP/allergen mouse model of severe allergic airway inflammation, which reproduces severe asthma and is steroid resistant, is almost completely dependent on PRMT5. PRMT5 was required for lung inflammation, airway remodeling and AHR, the clinical correlate of impaired breathing in human asthma. Remarkably, these effects were achieved just by eliminating PRMT5 from T cells while structural cells remained PRMT5 sufficient, indicating that PRMT5 in T cells drives multiple clinically-relevant aspects of this complex disease. Mechanistically, our data show that PRMT5-sufficient T cells are essential for pulmonary cholesterol metabolism, Th17 responses, and mixed granulocytic infiltration. Overall, our work demonstrates the importance of T cell PRMT5 in severe allergic lung inflammation and has potential implications for the pathogenesis and therapeutic targeting of severe asthma.

## Methods

### Allergen Mouse Model

Mouse studies were approved by The Ohio State University Institutional Animal Care and Use Committee under protocol #2019A00000108 (MG) and #2020A00000037 (JE). Homozygous *Prmt5*^*tm2c(EUCOMM)wtsi*^ mice were obtained and genotyped as described (15) and were provided food and water *ad libitum*. Litters were intranasally challenged starting at postnatal day 3 with PBS or mixed allergens (MA) (10µg *Alternaria Alternata*, 10µg *Aspergillus Fumigatus*, 10µg *Dermatophagoides Pteronyssinus* (all from Stallergenes Greer, Lenoir, NC), 10µg OVA plus 0.5µg c-di-GMP (both from InvivoGen, San Diego, CA)) 3 times per week for 7 weeks, no significant differences between genders(16).

### Bronchoalveolar lavage harvesting

Bronchoalveolar lavage fluid (BAL) was collected, processed, and stored for total and differential cell counts, cytokine, and sterol analyses as described (17). Total counts were performed using a hemocytometer. Cytospins were generated and differential cell counts were performed as described (18) using a modified Wright-Giemsa Stain (Newcomer Supply, 9112B, Middleton, WI).

### ELISA

Mouse CCL11, CXCL10, IL-4, IL-5, IL-13, IL-17A, IFNγ, and TNFα levels were measured in BAL supernatant using Meso Scale U-Plex assay (Meso Scale Diagonostics, Rockville, MD) following the manufacturer’s instructions. Plates were read on the MESO QuickPlex SQ 120 (Meso Scale Diagnostics). CXCL1 was measured in BAL using the mouse CXCL1 Quantikine ELISA kit (R&D Systems, Minneapolis, MN) following manufacturer’s instructions.

### Flow Cytometry

Single cell suspensions for flow cytometry were prepared as described (19) and stained with the following antibodies: anti-CD45-FITC (30-F11), anti-CD3-BV421 (17A2), anti-CD4-PE (A161A1), anti-IFNγ-Alexa700 (XMG1.2), anti-IL-4-APC (11B11), and anti-IL-17A-APC (TC11-18H10.1) (all from Biolegend). Data were acquired with a BD LSRII flow cytometer (BD Biosciences, San Diego, CA) and analyzed with FlowJo.

### Histological Analyses

Left lung lobes were inflated with 10% neutral buffered formalin at 25 cm H_2_O and paraffin embedded. Slides were stained with H&E to assess immune cell infiltration/aggregation in peribronchial and perivascular spaces (0-4 scale) and inducible bronchial-associated lymphoid tissue (iBALT) formation. Slides were stained with Alcian Blue-Periodic Acid Schiff (PAS) to quantify mucous cells in the airway epithelium. Photomicrographs of four different airways were taken at 100X on Lionheart microscope (Biotek, Winooski, VT). PAS positive and negative cells were quantified blinded using ImageJ (NIH) and reported as a percentage of total airway epithelial cells counted.

### Immunohistochemistry

Formalin-fixed, paraffin-embedded left lung lobe 6µm sections were used for immunohistochemical staining for α-Smooth Muscle Actin (α-SMA). Sections were deparaffinzed with xylene and rehydrated with graded ethanol. Antigen retrieval was with 10mM sodium citrate at 100°C for 1h. Blocking: 4% goat serum/0.04% Triton X-100/TBS; primary antibody: mAb anti-α-SMA at 1:100 (Millipore Sigma, A2547); secondary antibody: anti-mouse-Alexa 594 at 1:1000 (Millipore Sigma, F0257). Images were taken at 100X magnification on Lionheart (Biotek).α-SMA area was analyzed using ImageJ (NIH). Measured airway smooth muscle (ASM) mass was normalized to length of airway basement membrane and reported as ASM mass (µm^2^ per µm basement membrane).

### Lung Function

The exposed trachea of an anesthetized mouse was cannulated with a 19-gauge blunt-tip cannula. While attached to Y-tubing on the Flexivent (SCIREQ, Montreal, Quebec, Canada), we assessed airway hyperresponsiveness (AHR) by performing Snapshot (resistance, compliance, elastance) maneuvers after nebulization with PBS and a methacholine dose response (Millipore-Sigma, St. Louis, MO). AHR is reported as total resistance in response to methacholine.

### Myeloperoxidase assay

The tissue isolation and the myeloperoxidase assay were performed as described (20). Briefly, diluted sample MPO was bound to anti-MPO coated plates (Hycult Biotech HK210-01) followed by washes. Fluorescence (ex 535nm, em 590nm) after addition of H_2_O_2_ and ADHP was acquired in kinetic mode (10min, every 30 sec) on a Biotek Cytation 1. Results reported as RFU/second.

### Western blotting

Right lung lobes were homogenized and lysed in RIPA buffer (10 mM Tris pH 8.0, 5 M NaCl, 0.5 M EDTA, 1% Triton X-100, 0.1% SDS, 0.1% sodium deoxycholate) containing protease and phosphatase inhibitors (ThermoFisher Scientific). Primary antibodies: PRMT5 (Abcam ab31751, 1:1000), H4R3 (MilliporeSigma SAB4300870, 1:500), SYM10 (MilliporeSigma 07-412, 1:300), ROR-γt (Life Technologies 14-6981-82, 1µg/ml) and β-actin (Sigma-Aldrich A1978, 1:50,000). Secondary antibodies (1:20,000): donkey anti-rabbit or anti-rat 800CW and donkey-anti-mouse 680RD (Li-cor). Blots were imaged on an Odyssey-CLx and quantified with Image Studio software (Li-Cor).

### Mass Spectrometry of cholesterol and cholesterol precursors in lung infiltrating cells and BAL

BAL cells were pelleted by centrifugation and cell-free supernatants collected. Lungs were processed as described in the Flow Cytometry section followed by a 70-30% isotonic Percoll gradient and pelleted. Both were stored at −80 until mass spectrometry processing. 300μL BAL or gradient-isolated lung infiltrating cell pellets were spiked with sterol internal standards and processed for derivatization and LC–MS/MS analysis of sterols as described (21). Endogenous sterols were quantified by using the matching deuterated sterols and reported nmol/mL for BAL or nmol/million for cells or the ratio to cholesterol. Full list of SRM of sterols was reported previously (21). Sterol numbers were calculated relative to ml of BAL (for BAL) or total number of infiltrating cells (for lung infiltrating cells) and then normalized to the cholesterol/ml or cholesterol/cells value for that sample.

### Real-Time PCR

300ng of RNA isolated from right lungs using Zymo Direct-zol RNA Isolation Kit (Zymo Research, Irvine, CA) was transcribed with Oligo dT_12-18_ primers and SuperScript IV (ThermoFisher Scientific). Quantitative real-time PCR with TaqMan gene expression assays (ThermoFisher Scientific, all 4331182; HPRT Mm00446968_m1, PRMT1 Mm00480135_g1, PRMT2 Mm01173299_m1, PRMT3 Mm00659701_m1, Carm1 Mm00491417_m1, PRMT5 Mm00550472_m1, PRMT6 Mm01206465_s1, PRMT7 Mm00520495_m1, PRMT8 Mm01182914_m1, PRMT10 Mm00626834_m1) were run on a QuantStudio 3 96-well Real-Time PCR system (ThermoFisher Scientific). Results were analyzed by the comparative Ct method.

### Statistical analyses

Statistical analyses were performed using GraphPad Prism. Student’s t test or 1-way ANOVA followed by Sidak’s post-hoc multiple comparisons test was used as appropriate. Pearson correlation was used for correlation analyses.

## Results

### PRMT5 protein and its symmetric dimethylation activity are increased during severe lung inflammation

We have recently developed a model where chronic GMP and mixed allergen exposure results in chronic corticosteroid-resistant lung inflammation, airway remodeling and AHR (16). Since PRMT expression has been observed to be closely regulated at the protein level rather than the transcript level (22), we evaluated the protein expression of the major Type I and II PRMTs, namely PRMT1 and PRMT5, and their methylation marks in lung tissue from 7-week PBS or GMP/MA-exposed mice (**Fig. 1A**). Type I PRMTs catalyze asymmetric dimethylation (ADM) and type II PRMTs catalyze symmetric dimethylation (SDM). We found that both PRMT1 and PRMT5 were induced in the lung of GMP/MA vs. PBS-exposed mice for 7 weeks (**Fig. 1 B-C**). In contrast to protein induction, both PRMT1 and PRMT5 were significantly decreased at the transcript level (**Fig. S1A-B**), suggesting these PRMTs are regulated at the protein level during lung inflammation. Consistent with induced PRMT1 and PRMT5 protein in the lung, both SDM (indicative of PRMT5 activity) and ADM (indicative of PRMT1 activity) were increased in allergic lung tissue, with robust detection of SDM (**Fig. 1 D-E**). PRMT induction may originate from a number of changes in the lung, from changes in expression in resident cells to newly infiltrating immune cells. PRMT5 is expressed in T cells where we have shown that it promotes Th17 differentiation and IL-17A production (15). In turn, IL-17A is linked to neutrophilic inflammation. To identify potential links between lung PRMT5 induction/activity to Th17 inflammation, we performed correlation analyses. Lung PRMT5 protein and its SDM mark were significantly and positively correlated to bronchoalveolar lavage (BAL) IL-17A (**Fig. 1F-G**). Overall, these findings suggest that the type II methyltransferase PRMT5 contributes to severe lung inflammation characterized by Th17 inflammation and neutrophilic and eosinophilic (T2/T17) components.

**Figure 1.**
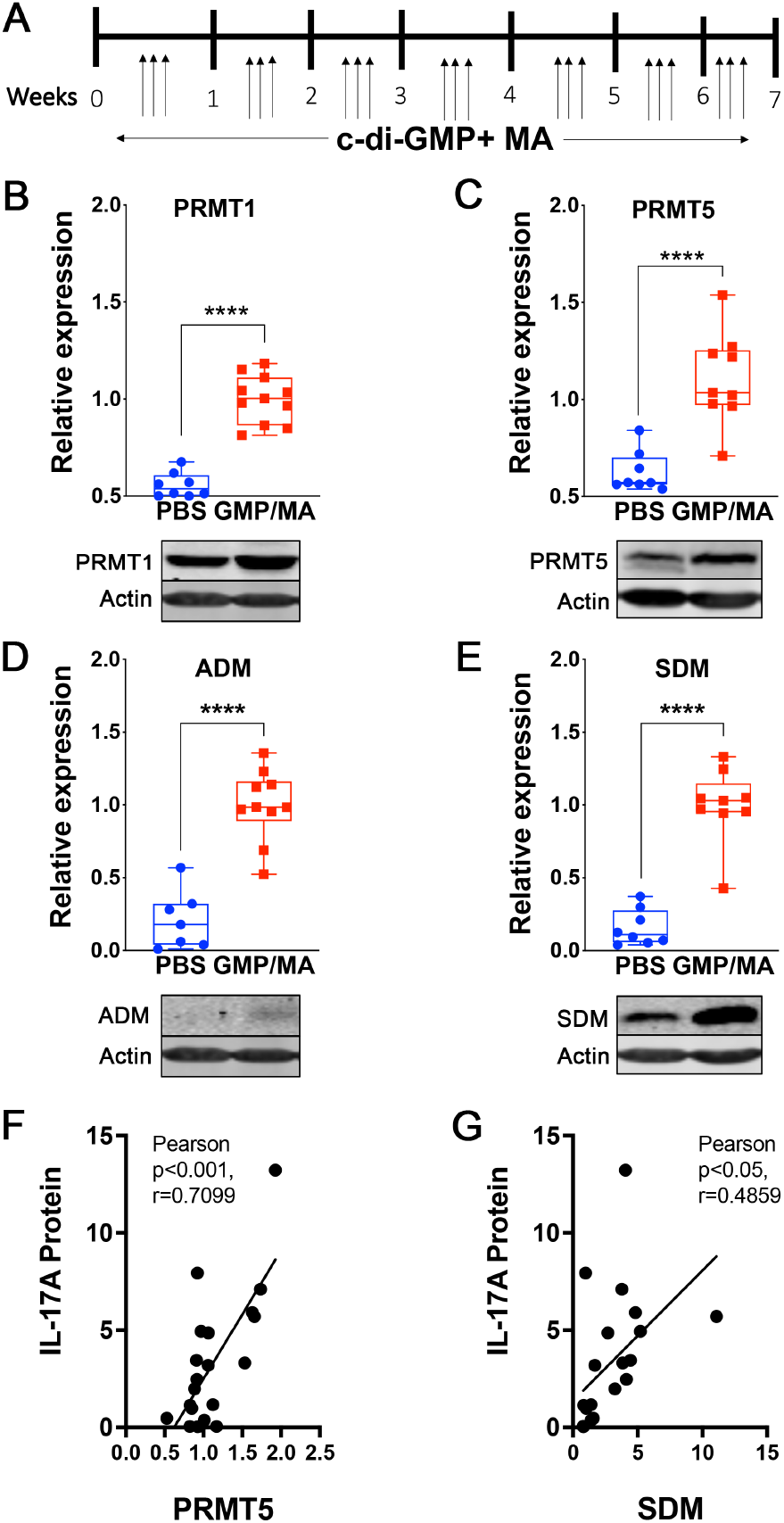
PRMT5 expression and activity are increased during severe airway inflammation and positively correlate with IL-17 responses. (A) Schematic of the GMP/MA allergic airway inflammation model: C57Bl6/J mice are exposed to c-di-GMP and mixed allergen (MA: 10 ug each of *Aspergillus fumigatus, Alternaria alternata*, house dust mite and ovalbumin) 3x/week for 7 weeks and analyses are performed at 7 weeks. (B-E) Representative blot and quantification of PRMT1 (B), PRMT5 (C) protein expression, PRMT1’s asymmetric dimethylation (ADM) mark (D) and PRMT5’s symmetric dymethylation (SDM) mark (E), all evaluated by Western blot, in whole lung lysate (n= 6-9/group, pooled from two independent experiments, ^****^p<0.0001, unpaired t test). (F-G) Pearson correlation analysis of bronchoalvolar lavage (BAL) IL-17 and either PRMT5 (F) or its methylation mark (G). Data from n= 21, pooled from two independent experiments, stats. ^*^p<0.5, ^***^p<0.001.

### PRMT5 Promotes Airway Inflammation and Remodeling

Given the increases in PRMT5 expression and its importance in Th17 differentiation, we examined the role of T cell PRMT5 in the GMP/MA model, which exhibits structural and functional changes that are insensitive to steroids (16). To evaluate the role of PRMT5 in T cells, we exposed mice with a CD4-cre driven PRMT5 deletion in T cells (PRMT5^fl/fl^CD4-Cre^+^ mice subsequently referred to as KO) or appropriate littermate controls (PRMT5^fl/fl^CD4-Cre^-^ mice, subsequently referred to as WT in figures) to GMP/MA (15). We have validated efficient and cell specific deletion in this model (see (15)). Quantification of lung inflammation showed significant immune cell infiltration in the perivascular and peribronchiolar spaces in GMP/MA challenged WT mice (**Fig. 2A-B**). These mice also developed inducible broncho-associated lymphoid tissue (iBALTs) (**Fig. 2A, C**), a pathological feature implicated in severe allergic airway inflammation (23–25). Immune cell infiltration and aggregation around airways and within lung tissue was significantly reduced in GMP/MA-challenged KO mice (**Fig. 2A-C**). Similarly, GMP/MA-induced mucous cell abundance and airway smooth muscle mass were significantly reduced in GMP/MA-challenged KO mice (**Fig. 2A, D-E**). Lack of airway remodeling and thickening in GMP/MA-exposed KO mice was accompanied by decreased AHR compared to WT GMP/MA-challenged mice (**Fig. 2F**). These findings demonstrate the importance of CD4 T cell PRMT5 in the structural and functional changes during severe allergic airway inflammation.

**Figure 2.**
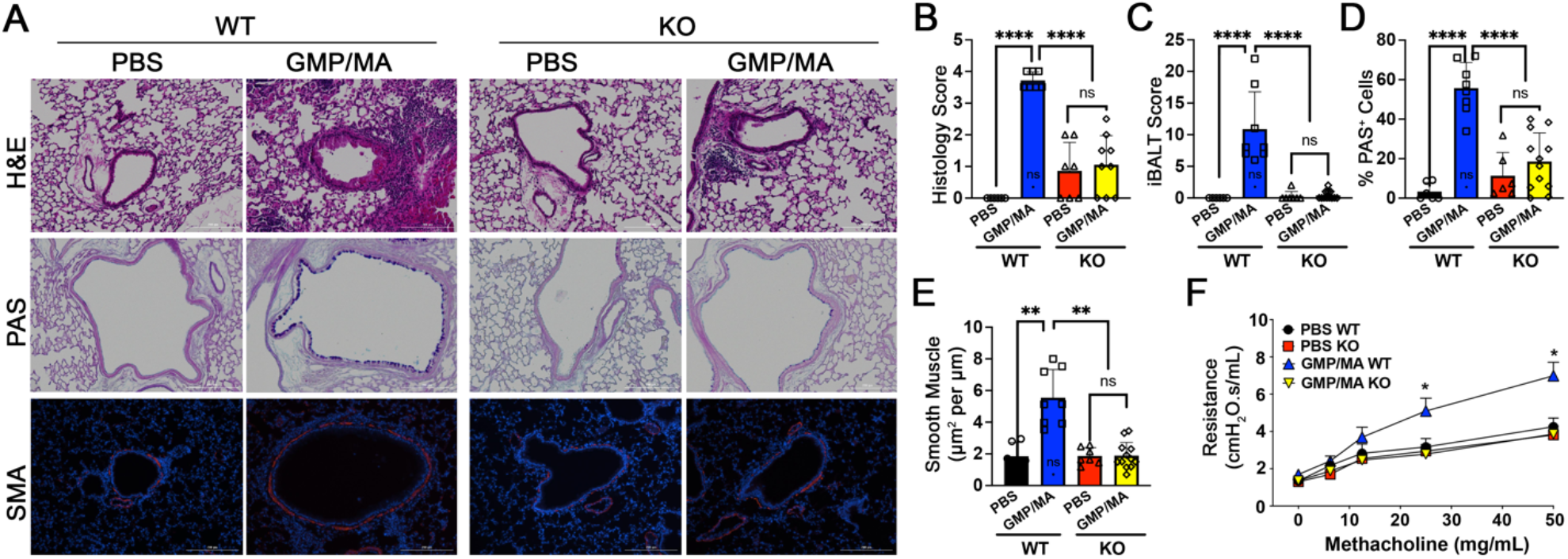
PRMT5 is key for development of prominent pathological features and airway hyperresponsiveness (AHR). Wild-type control (T-PRMT5^fl/fl^) and CD4 PRMT5 KO (T-PRMT5^fl/fl^CD4-cre) mice were exposed to GMP/MA for 7 weeks and evaluated for lung pathology and AHR. (A-B) Lung inflammation and (C) iBALT formation are reduced in CD4 PRMT5 KO mice. Structural changes including (D) mucous cell abundance and (E) airway smooth muscle mass are reduced in CD4 PRMT5 KO mice. (F) AHR fails to increase in CD4 PRMT5 KO mice challenged with GMP/MA. Data are presented as mean ± SE, n=6-12/group. ^*^ p<0.05.

### PRMT5 in T cells promotes mixed granulocytic lung infiltration with eosinophils and neutrophils

To explore the immunological mechanisms associated with the robust improvements in pathology and AHR observed in GMP/MA-challenged KO mice, we evaluated immune infiltrates in the BAL. At 7 weeks, WT mice exposed to GMP/MA developed mixed granulocytic inflammation characteristic of the model, while similarly exposed KO mice showed an almost complete reduction of BAL eosinophils and neutrophils, as well as macrophages (**Fig. 3A**). Neutrophils contribute to lung pathogenesis through release of granules that contain oxidative enzyme activity.

**Figure 3.**
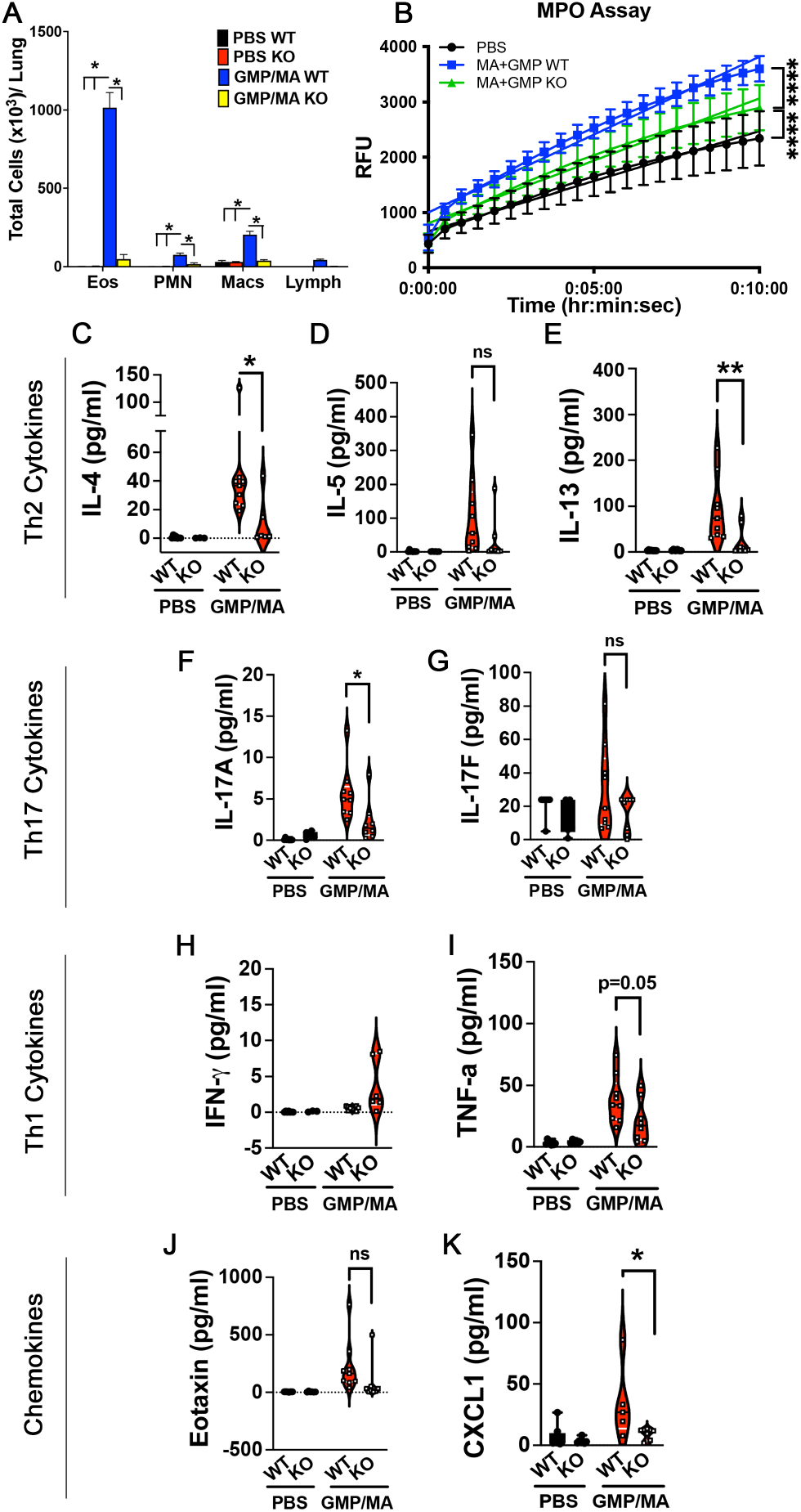
PRMT5^fl/fl^CD4-cre^+^ (KO) and PRMT5^fl/fl^CD4-cre^-^ (corresponding functional wild-type control, labeled WT) mice were exposed to GMP/MA for 7 weeks and evaluated for (A) BAL infiltrating cells (n= 7-9, pooled from >/= 3 independent experiments, ^*^ p<0.05, ANOVA followed by Tukey’s multiple comparison), (B) lung neutrophil myeloperoxidase activity (n=9-15/group, pooled from >/= 3 independent experiments, ^****^ p<0.0001 ANOVA followed by Tukey’s multiple comparison of slopes). (C-K) BAL cytokines and chemokine levels are reduced in KO mice. T2 cytokines: (C) IL-4, (D) IL-5, (E) IL-13; T17 cytokines: (F) IL-17A and (G) IL-17F; T1 cytokines (H) IFN- and (I) TNF-; Chemokines: (J) Eotaxin and (K) CXCL1. Data are presented as mean ± SE, n=8-9/group. ^*^ p<0.05.

Therefore, we evaluated neutrophil myeloperoxidase (MPO) activity, which has been shown to serve as a biomarker of neutrophil infiltration in lung tissue. We found that MPO activity was substantially decreased in the lung parenchymal tissue of KO mice (**Fig. 3B**). Eosinophil and neutrophil infiltration are promoted by cytokines and chemokines associated with T2 and T17 immune responses, respectively. We found that BAL IL-4, IL-13, and IL-17A were significantly reduced in KO mice, whereas IL-5 and IL-17F did not reach significance (**Fig. 3C-G**). For T1-associated cytokines, BAL IFN-γ levels were increased in GMP/MA KO mice and TNF-was significantly reduced in KO mice (**Fig. 3H-I**). For neutrophil and eosinophil chemoattractants, BAL CXCL1 and CCL11 levels were reduced in KO mice (**Fig J-K**).

### PRMT5 deletion in T cells suppresses desmosterol content and Th17 responses in lung infiltrating cells

The observed decreases in T2 and T17 cytokines suggest that Th2 and Th17 responses that promote neutrophil and eosinophil infiltration are impaired after PRMT5 deletion in T cells. To address this, we first evaluated whether loss of PRMT5 on T cells had a non-specific effect on lung T cell loss. We found that naïve KO mice had normal lung total CD4 T cell numbers (WT: mean 2.5×10^5^; KO: 5×10^5^; **supplemental Fig. 2A**). In addition, PBS and GMP/MA-exposed KO mice had normal naïve Th cell numbers (**supplemental Fig. 2B**). In contrast, we did observe a decrease in lung memory T cells in KO mice exposed to GMP/MA (**supplemental Fig. 2C**). While Th1 and Th2 cell % remained increased in the lung, Th17 cell % in GMP/MA KO was comparable to PBS mice (**Fig. 4A**). Total cell numbers of Th1 and Th2 cells were not significantly decreased while Th17 cells were robustly suppressed (**Fig. 4B**). Taken together, these data suggest PRMT5 does not directly promote Th1 or Th2 cells, but rather, Th17 cell populations. Our previous work showing PRMT5 deficiency does not impair *in vitro* Th2 differentiation of isolated naïve CD4 Th cells further support this idea (15). We then performed sterol analyses since CPI have been shown to drive ROR-γt activity and Th17 differentiation. Indeed, we found reduced levels of the CPI desmosterol in lung infiltrating cells and the BAL of GMP/MA-exposed KO mice (**Fig. 4C-D)**. This decrease was not due to overall reductions in infiltrating cells, as desmosterol levels were normalized to total cholesterol. Other CPI sterols, such as zymosterol, were altered in BAL, but not lung infiltrating cells (**Supplemental Figure 3A-C**). If PRMT5 promotes lung Th17 responses, we would expect increased lung ROR-γt expression. Accordingly, we observed a PRMT5-dependent increase in ROR-γt expression in the lung (**Fig. 4E**). Overall, these data suggest that loss of PRMT5 in T cells impairs biosynthesis of sterols that promote ROR-γt-dependent Th17 lung responses.

**Figure 4.**
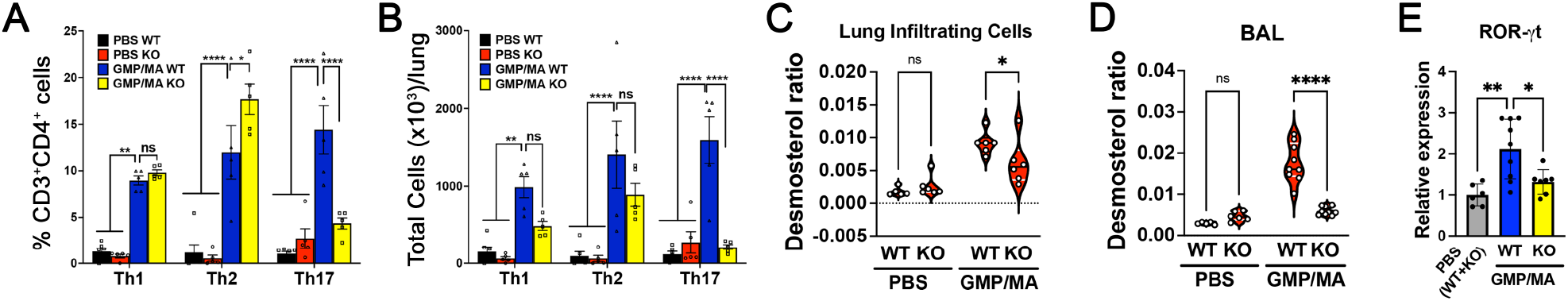
PRMT5^fl/fl^CD4-cre^+^ (KO) and PRMT5^fl/fl^CD4-cre^-^ (WT, corresponding functional wild-type control) mice were exposed to GMP/MA for 7 weeks and evaluated for (**A-B**) lung infiltrating Th1, Th2 and Th17 cell % (**A**) and cell number (**B**) by flow cytometry (n= 5-6, pooled from >/= 3 independent experiments, ^*^ p<0.05, ^**^ p<0.01, ^****^ p<0.0001, ANOVA followed by Tukey’s multiple comparison), (**C-D**) CPI desmosterol, normalized to total cholesterol, in lung infiltrating cells (**C**) and BAL (**D**) (n= 5-6, pooled from two independent experiments, ^*^ p<0.05, ANOVA followed by Sidak’s multiple comparison) and (**E**) ROR-γt protein expression in whole lung by Western blot. (n= 6-9, pooled from two independent experiments, ^*^ p<0.05, ^**^ p<0.01, ANOVA followed by Sidak’s multiple comparison test, ^*^ p<0.05.

## Discussion

In this manuscript, we identify a key role for PRMT5 as a driver of severe allergic airway inflammation. While both PRMT1 and PRMT5 were induced in the GMP/MA lung, we show that deletion of PRMT5 in T cells protects mice from severe lung inflammation with mixed granulocytic infiltrates, bronchial associated lymphoid tissue development, airway remodeling and AHR. In addition, we find that PRMT5 in T cells promotes lung inflammation rich in the CPI desmosterol, lung ROR-γt expression, and Th17 cells. Although Th2 cells were not significantly decreased in T-PRMT5 KO mice, both T2 (IL-4) and T17 (IL-17A) cytokines were decreased, as well as eosinophil and neutrophil infiltration.

Increased aginine methylation has been described in models of allergic airway inflammation, as well as in human asthma (12, 13). Initial work largely detected ADM during lung inflammation, although SDM has also been observed (12, 13). We found that the two major PRMTs catalyzing SDM, namely PRMT5, and ADM, namely PRMT1, are induced in lung tissue during GMP/MA inflammation. Both SDM and ADM were also increased and may in principle play a role in lung disease. Early work showed that the AMI-1 inhibitor initially described as a PRMT1 inhibitor can suppress airway inflammation (26). However, AMI-1 has since been shown to suppress PRMT5 as well, raising the possibility that targeting PRMT5 could mediate beneficial effects. Indeed, we find that sole deletion of PRMT5 in T cells is sufficient to halt multiple aspects of severe lung inflammation. It is important to note that, at least in T cells, PRMT5 has been found to regulate PRMT1 in a positive feedback loop (27–30). Therefore, part of PRMT5’s effects could be indirectly mediated by PRMT1 induction.

Loss of PRMT5 in T cells robustly decreased lung infiltrating Th17 cells in the GMP/MA model, with minor non-significant decreases in total Th1 and Th2 cells (and no decreases in percentages) also observed. We have recently found, using in vitro Th cell differentiation of isolated naïve Th cells, that PRMT5 is essential for murine Th17 differentiation (15). In those studies, we observed that the decreases in Th17 cell proportion observed in PRMT5 deficient Th cells were not caused by Th cell death, reduced proliferation, or gating artifacts (15). In addition, we found that PRMT5 deletion has less of a profound or no effect on Th1, Th2 and Treg differentiation (15). Overall, these results suggest a prominent role for PRMT5 in Th17 differentiation. The main cytokine produced by Th17 cells is IL-17A, a pro-neutrophilic cytokine that increases bone marrow neutrophil production, as well as neutrophil recruitment into tissues (31). Accordingly, we found reduced neutrophils in PRMT5 KO mice. Interestingly in addition to neutrophils, we observed significant reduction in eosinophil infiltration. This leads to the somewhat provocative idea that Th17 responses and/or IL-17A is an important driver of both eosinophilic and neutrophilic lung infiltration, at least in certain contexts. This idea is supported by observations in fungal allergen or mixed allergen models (32–34). Since human allergen exposures are often varied and combined, it is possible that these findings are highly significant to human asthma.

We hypothesize that the molecular mechanism by which PRMT5 in T cells regulates Th17 responses in lung inflammation is via modulation of cholesterol metabolism. Sterols are increasingly recognized as active lipid mediators of importance in immune and other biological processes. Several precursors to cholesterol, as well as cholesterol itself, have strong ROR-γt agonistic activity and drive Th17 differentiation (15, 35, 36). In particular, desmosterol is a highly active ROR-γt agonist and we found it increased in infiltrating cells from GMP/MA challenged mice, while its BAL and infiltrating cell levels were reduced in KO mice. Since we analyzed mononuclear cells isolated by Percoll gradient, T cells likely contribute to this desmosterol increase. We have previously observed T-cell specific decreases in cholesterol pathway biosynthesis enzymes *Lss, Cyp51a, Tm7sf2, Nsdhl, Msmo1* and *Sc5d* after PRMT5 deletion (15), as well as loss of ROR-γt activity in T cells after PRMT5 knockdown (15). ROR-γt activity was also decreased in GMP/MA lungs of T cell PRMT5 KO mice. Overall, our results and prior literature suggest that reductions in cellular desmosterol content during severe airway inflammation may promote Th17 responses in the lung.

In addition to the control of adaptive immune responses, cholesterol and fatty acid metabolism changes in innate immune cells have been shown to modulate their activity. TLR signaling in macrophages induces metabolic reprogramming in macrophages, with increased lanosterol and desmosterol levels (37, 38). These changes can dampen inflammatory signals while increasing the microbial activity of macrophages (37, 38). On the other hand, excess cholesterol levels in DCs and macrophages have been observed to drive NLPR3 inflammasome activation, production of pro-inflammatory cytokines such as IL-1β and lymphocyte activation (39, 40). However, deficiencies in serum proteins (i.e. apolipoproteins) and/or receptors known to facilitate cholesterol uptake have been associated with increased neutrophilia and Th17 responses in asthma (41–45). In humans, cholesterol targeting statin drugs are often considered to have anti-inflammatory effects. However, several statin trials in did not observe clinical asthma improvements (46). More recently, three large studies retrospective studies encompassing more than 100,000 patients showed a reduction in emergency visits for statin-treated asthmatic patients (47–49, 49), suggesting cholesterol metabolism may contribute to severe asthma where exacerbations and hospitalizations are frequent. Overall, these studies highlight the need of additional research to better understand the roles and mechanisms by which cholesterol metabolism and its therapeutic modulation play during immune responses and asthma.

In summary, our work demonstrates the importance of T cell PRMT5 in severe allergic lung inflammation and has potential implications for the pathogenesis and therapeutic targeting of severe asthma, particularly asthma with mixed eosinophilic and neutrophilic infiltration. Since small molecule PRMT5 inhibitors have been developed and are currently being evaluated in clinical trials, they may serve as therapeutic options for asthma patients and others with Th17-mediated diseases.

## Acknowledgements

This work was supported by funds from the the Drug Development Institute (MG), the NIH-NHLBI Center for Accelerated Innovations-Cleveland Clinic (NCAI-CC, MG), the School of Health and Rehabilitation Sciences and the Division of Medical Laboratory Science (MG), OSU College of Medicine and Nationwide Children’s Hospital Collaborative Pilot Grant (MG and RB) and NIH National Institute of Allergy and Infectious Diseases grants R01AI121405 and 1R21AI127354 (MG), the Comprehensive Cancer Center Genetically Engineered Mouse Modeling Core (Core Cancer Center Support Grant P30CA016058), National Institutes of Health R00 HL131682 (RB), R01 HL155095 (RB), startup funds from the Abigail Wexner Research Institute at Nationwide Children’s Hospital (RB). The content is solely the responsibility of the authors and does not necessarily represent the official views of the NIH.

