## Supplemental Figures 1-3 for "PRMT5 in T cells drives Th17 responses, mixed granulocytic inflammation and severe allergic airway inflammation"

**Supplemental Data**

Supplemental Figure 1

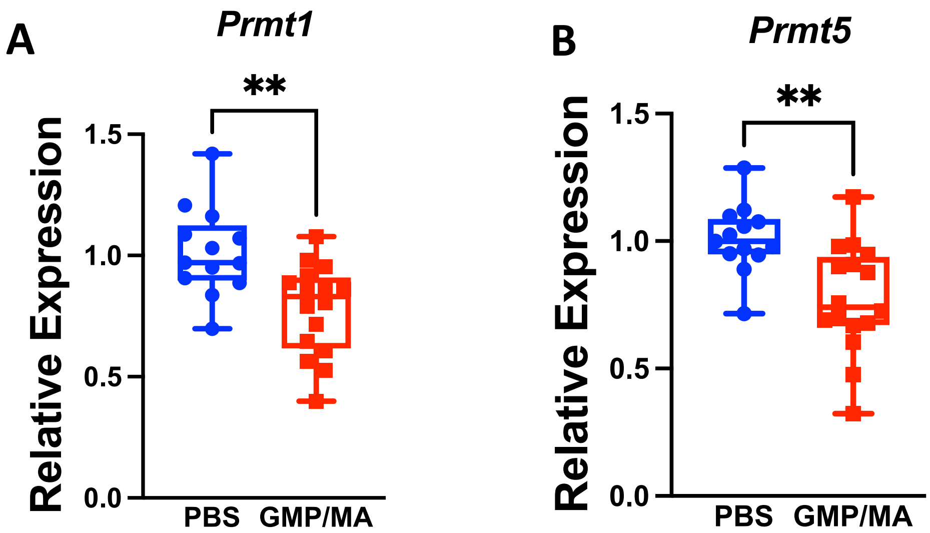


**Supplemental Figure 1.** *Prmt1* and *Prmt5* transcripts are decreased in the lung during GMP/MA lung inflammation. Wild-type C57Bl6/J mice were exposed to GMP/MA for 7 weeks, lungs harvested and processed for RNA isolation and Real-Time PCR quantification of *Prmt1* (A) and *Prmt5* (B) transcript abundance. N=13-16, pooled from two independent experiments, ** p<0.01, t-test.

Supplemental Figure 2

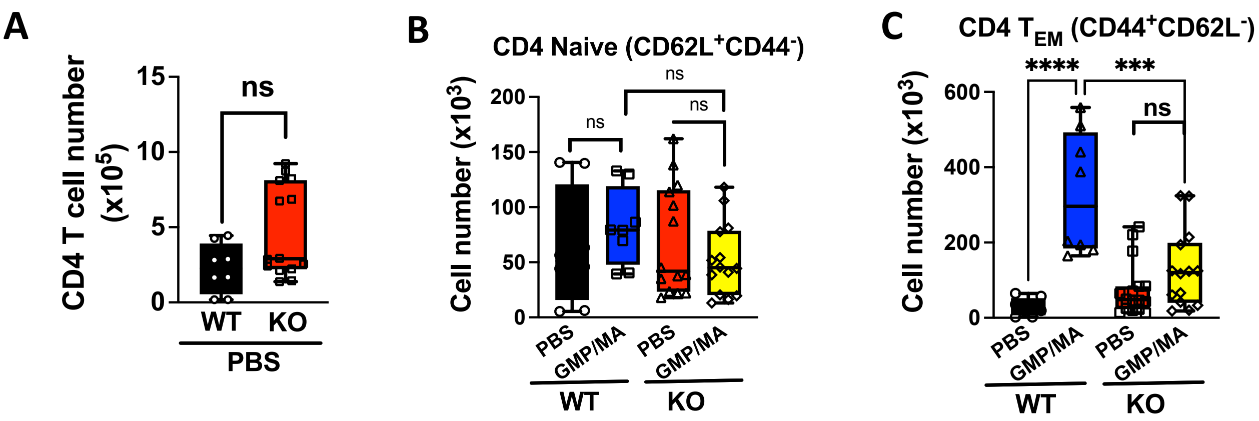


**Supplemental Figure 2.** KO mice harbor normal lung total and naïve T cell numbers but show lung TEM impairments during GMP/MA lung inflammation. KO and corresponding functional wild-type control mice were exposed to GMP/MA for 7 weeks and evaluated for lung parenchyma (**A**) total CD4 T cells, (B) naïve CD4 T cells or (**C**) T effector memory (TEM) cells. T test (A) or ANOVA followed by multiple comparison Sidak's test (B-C), **** $p<0.001$. Data from two experiments with n=4-6 biological replicates/group.

Supplemental Figure 3

A

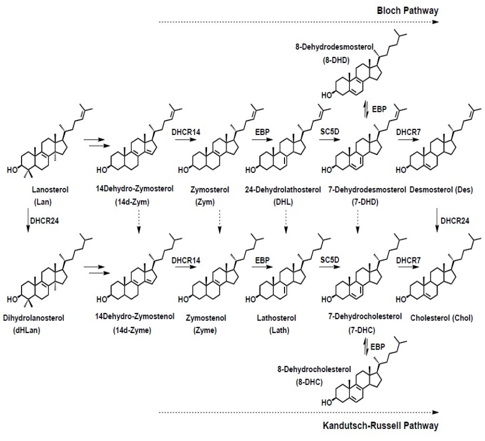


B

| | | | | | GMP/MA (WT vs KO) |
|---|---|---|---|---|---|
| BAL | PBS WT | PBS KO | GMP/MA WT | GMP/MA KO | p value (ANOVA Sidak) |
| Lan | 1.75E-04 +/- 7.27E-05 | 1.68E-04 +/- 8.61E-05 | 1.01E-03 +/- 3.89 E-04 | 2.66E-04 +/- 2.38E-04 | **** |
| 14dZym | 2.6E-05 +/- 6.36E-06 | 5.94E-05 +/- 9.78E-05 | 2.02E-04 +/- 6.95E-05 | 2.95E-05 +/- 1.79E-05 | **** |
| Zym | 3.0E-04 +/- 5.94E-05 | 3.60E-05 +/- 9.69E-05 | 8.37E-04 +/- 1.49E-04 | 5.38E-04 +/- 1.18E-04 | **** |
| DHL | 1.08E-03 +/- 1.78E-04 | 1.41E-03 +/- 3.18E-04 | 3.69E-03 +/- 7.16E-04 | 1.87E-03 +/- 2.46E-04 | **** |
| 7DHD | 1.03E-03 +/- 1.22E-04 | 1.50E-03 +/- 3.51E-04 | 1.61E-03 +/- 3.17E-04 | 1.84E-03 +/- 2.14E-04 | ns |
| 8DHD | 8.70E-05 +/- 1.26E-05 | 1.63E-04 +/-2.34E-05 | 1.93E-04 +/- 4.21E-05 | 1.81E-04 +/- 5.31E-05 | ns |
| Des | 2.99E-03 +/- 3.15E-04 | 4.69E-03 +/- 1.14E-03 | 1.77E-02 +/- 4.71E-03 | 6.00E-03 +/- 1.02E-03 | **** |
| dHLan | ND | 4.14E-05 +/- 6.15E-05 | 6.26E-05 +/- 5.18E-05 | 1.05E-04 +/- 2.58E-05 | ns |
| 14dZyme | 9.37E-04 +/- 1.06E-04 | 1.24E-03 +/- 3.07E-04 | 4.72E-03 +/- 1.28E-03 | 1.58E-03 +/- 2.86E-04 | **** |
| Zyme | ND | ND | ND | ND | **** |
| Lath | 1.09E-03 +/- 1.42E-04 | 1.30E-04 +/- 2.08E-04 | 2.74E-03 +/- 4.03E-04 | 1.85E-03 +/- 3.84E-04 | **** |
| 7DHC | 2.6E-03 +/- 1.08E-03 | 2.03E-03 +/- 2.03E-03 | 2.20E-03 +/- 9.38E-04 | 2.5E-03 +/- 1.64E-03 | ns |
| 8DHC | 2.3E-03 +/- 1.77E-04 | 2.07E-03 +/- 3.46E-04 | 2.05E-03 +/- 2.14E-04 | 2.2E-03 +/- 2.74E-04 | ns |

C

| | | | | | GMP/MA (WT vs KO) |
|---|---|---|---|---|---|
| Infiltrating cells | PBS WT | PBS KO | GMP/MA WT | GMP/MA KO | p value (ANOVA Sidak) |
| Lan | 5.72E-05 +/-5.43E-05 | 1.02E-04 +/- 8.54E-05 | 3.11E-04 +/- 1.21E-04 | 2.02E-04 +/- 9.67E-05 | ns |
| 14dZym | 1.29E-05 +/- 9.26E-06 | 2.12E-05 +/- 1.16E005 | 4.49E-05 +/- 3.00E-05 | 5.85E-05 +/- 3.25E-05 | ns |
| Zym | 1.56E-04 +/- 4.91E-05 | 1.85E-04 +/- 8.25E-05 | 3.70E-04 +/- 7.94E-05 | 4.78E-04 +/- 2.40E-04 | ns |
| DHL | 3.74E-04 +/- 7.06E-05 | 5.13E-04 +/- 2.25E-04 | 1.47E-03 +/- 2.06E-04 | 1.64E-03 +/- 9.85E-04 | ns |
| 7DHD | 3.61E-04 +/- 9.81E-05 | 5.12E-04 +/- 2.52E-04 | 1.26E-03 +/- 3.26E-04 | 1.34E-03 +/- 6.16E-04 | ns |
| 8DHD | 6.65E-05 +/- 5.30E-05 | 4.48E-05 +/- 2.73E-05 | 9.27E-05 +/- 2.21E-05 | 7.54E-05 +/- 2.28E-05 | * |
| Des | 1.79E-03 +/- 6.16E-04 | 2.64E-03 +/- 1.56E-03 | 9.20E-03 +/- 1.72E-03 | 6.16E-03 +/- 3.44E-03 | ns |
| dHLan | 0.00E+00 +/- 00E+00 | 1.25E-06 +/- 2.49E-06 | 0.00E+00 +/- E00+00 | 0.00E+00 +/- 00E+00 | ns |
| 14dZyme | 3.96E-04 +/- 1.57E-04 | 6.01E-04 +/- 3.43E-04 | 2.41E-03 +/- 5.40E-04 | 1.62E-03 +/- 9.97E-04 | ns |
| Zyme | ND | ND | ND | ND | ns |
| Lath | 7.14E-04 +/- 1.52E-04 | 9.91E-04 +/- 2.79E-04 | 1.42E-03 +/- 3.38E-04 | 4.98E-03 +/- 8.90E-03 | ns |
| 7DHC | 2.78E-03 +/- 3.85E-03 | 1.11E-03 +/- 7.49E-04 | 1.58E-03 +/- 6.23E-04 | 2.03E-03 +/- 1.24E-03 | ns |
| 8DHC | 1.19E-03 +/- 2.65E-04 | 1.25E-03 +/- 1.08E-04 | 1.50E-03 +/- 1.42E-04 | 1.38E-03 +/- 2.70E-04 | ns |

**Supplemental Figure 3. (A)** Bloch and Kandutsch-Russell pathways of cholesterol biosynthesis. (B-C) Cholesterol pathway intermediate levels detected in (B) BAL or (C) gradient-isolated lung infiltratting mononuclear cells in WT and KO mice exposed to either PBS or GMP/MA for 7 weeks. Cholesterol pathway intermediate values are shown normalized for (A) BAL volume or (C) total number of infiltrating cells collected, followed by normalizing to total cholesterol content. (n= 5-6, pooled from two independent experiments, * $p<0.05$, ANOVA followed by Sidak's multiple comparison).
